# Non-rainfall moisture: a key driver of carbon flux from standing litter in arid, semiarid, and mesic grasslands

**DOI:** 10.1101/696666

**Authors:** Sarah E. Evans, Katherine E. O. Todd-Brown, Kathryn Jacobson, Peter Jacobson

**Affiliations:** Kellogg Biological Station, Ecology, Evolutionary Biology, and Behavior Program, Department of Integrative Biology, Department of Microbiology and Molecular Genetics, Michigan State University, Hickory Corners, MI, USA; Pacific Northwest National Laboratory, Richland, WA, USA; Wilfred Laurier University, Waterloo, Ontario, Canada; Department of Biology, Grinnell College, Grinnell, IA, USA

**Keywords:** fog, dew, non-rainfall moisture (NRM), standing litter, microbial decomposition, drylands, semiarid, mesic, modeling

## Abstract

Models assume that rainfall is the major source of moisture driving decomposition. Non-rainfall moisture (NRM: high humidity, dew, and fog) can also induce standing litter decomposition, but there have been few standard measurements of NRM-mediated decompositions across sites, and no efforts to extrapolate the contribution of NRM to larger scales to assess whether this mechanism can improve model predictions. Here we show that NRM is an important, year-round source of moisture in grassland sites with contrasting moisture regimes using field measurements and modeling. We first characterized NRM frequency and measured NRM-mediated decomposition in sites on the extreme dry and wet end of grassland systems: at two sites in the Namib Desert, Namibia (hyperarid desert) and at one site in Iowa, USA (tallgrass prairie). NRM was frequent at all sites (85-99% of hours that litter was likely to be wet were attributed to NRM) and tended to occur in cool, high-humidity periods for several hours or more at a time. NRM also caused respiration of standing litter at all sites when litter became sufficiently wet (>5% for fine litter and >13% for coarse), and contributed to mass loss, even in the Namib West site that had almost no rain. When we modeled annual mass loss induced by NRM and rain, and extrapolated our characterization of NRM decomposition to a final site with intermediate rainfall (Sevilleta, New Mexico, semiarid grassland), we found that models driven by rainfall alone underestimated mass loss, while including NRM produced estimates within the range of observed mass loss. Together these findings suggest that NRM is an important missing component in quantitative and conceptual models of litter decomposition, but there is nuance involved in modeling NRM at larger scales. Specifically, temperature and physical features of the substrate emerge as factors that affect the common microbial response to litter wetting under NRM across grasslands sites, and require further study. Hourly humidity can provide an adequate proxy of NRM frequency, but site-specific calibration with litter wetness is needed to accurately attribute decomposition to periods when NRM wets litter. Greater recognition of NRM-driven decomposition and its interaction with other processes (e.g. photodegradation) is needed, especially since fog, dew, and humidity are likely to shift under future climates.

**Manuscript highlights:**

- Non-rainfall moisture (NRM; humidity, fog, dew) induces decomposition in grasslands
- NRM decomposition depends on substrate type, and occurs at colder times than rain
- Including NRM (instead of rain alone) improved predictions of litter decomposition

## Introduction

Decomposition of plant litter and soil organic matter adds more CO_2_ to the atmosphere than fossil fuel use (Schlesinger and Andrews 2000). Thus, relatively small changes in decomposition will have large impacts on atmospheric CO_2_ concentrations and carbon-climate feedbacks. Despite this importance, our understanding of decomposition, and ability to predict how it will change under future climates, is limited. In particular, models, most of which use rainfall and temperature as the major climatic drivers of decomposition, consistently underestimate litter decay rates in drylands (Whitford and others 1981; Throop and Archer 2008), suggesting that mechanisms relevant to decomposition in these areas are omitted. Indeed, recent studies show that previously unrecognized processes such as photodegradation and soil-litter-mixing drive significant surface litter decomposition (Austin and Vivanco 2006; Gallo and others 2006; Throop and Archer 2008; Barnes and others 2011; King and others 2012; Baker and others 2015; Lin and others 2018).

An additional phenomenon that may explain underestimation of decomposition in drylands – and potentially other systems – is the stimulation of decomposition by non-rainfall moisture (NRM), or fog, dew, and high humidity. In semi-arid Mediterranean grasslands, Dirks et al., (2010) estimated that decomposition in the absence of both rain and photodegradation accounted for an 18% reduction in litter mass, which constituted up to 50% of annual decomposition in this system. They did not directly measure the effect of NRM on decomposition, but hypothesized that the decomposition they observed in rainless periods was due to atmospheric water vapor. Gliksman et al. (2016) quantified the influence of NRM-mediated decomposition (hereafter ‘NRM decomposition’) on mass loss at semiarid sites by manipulating microclimate, and saw a significant decrease in mass loss in litter bags when NRM and UV were excluded. The role of NRM in decomposition may extend beyond water-limited areas as well (Newell and others 1985; Kuehn and others 2004). For instance in wetlands, Kuehn et al. (2004) observed diel mineralization cycles of standing litter during rainless periods that corresponded with nightly dew formation, with carbon flux comparable to that emitted from soils and sediments.

Despite accumulating evidence that attests to the potential importance of NRM as a driver of decomposition, there have been few attempts to generalize the processes that control NRM decomposition across biomes, or scale NRM decomposition across space and time. Before NRM can be incorporated into conceptual and quantitative models, we need to know more about controls on NRM decomposition, and the best approaches for characterizing NRM frequency and duration in different ecosystems. Studies examining mechanistic controls on NRM decomposition, many performed in the laboratory, have highlighted several underlying drivers of NRM decomposition. Dirks et al., (2010) directly measured litter absorption of atmospheric water vapor in the laboratory, and saw stimulation of microbial activity. We showed that litter from the Namib Desert exhibited significant C flux under simulated nighttime dew and fog (Jacobson and others 2015), beginning within 5 minutes after gravimetric moisture exceeded a critical threshold, and lasting for 10 hours (as long as litter was wet). We also found that substrate type may be an important control on NRM decomposition; short periods (2 hours) of high relative humidity (>95%) induced microbial respiration, but only in fine-textured litter (e.g. grass leaves) and not in coarse tillers (stems) (Jacobson and others 2015). Further, litter position affects NRM decomposition – standing litter becomes wetter with nighttime humidity and has higher mass loss than surface litter (Almagro and others 2015; Wang and others 2017a; Gliksman and others 2018) – highlighting the importance of position on measurements of both NRM frequency (Sentelhas and others 2008) and litter decomposition.

Equally important as a mechanistic understanding is a quantification of the impact of this phenomenon at regional and annual scales. Few attempts have been made to characterize NRM across biomes, and even fewer to extrapolate its contribution to (e.g. annual) C loss. This is in contrast to the vast efforts made to monitor rainfall frequency and understand the effect of rainfall on soil moisture and C flux. Climatic variables that result in NRM, like humidity and diel temperature cycles, are fundamentally different from those affecting rainfall (McHugh and others 2015), and direct measurements of condensed water resulting from NRM, such as leaf wetness sensor measurements, are rarely included in standard meteorological measurements (Uclés and others 2015), or collected while monitoring litter respiration or mass loss. Further, measurements of humidity are typically made at standard height of 1.5 m, rather than at lower elevations representative of litter position, where RH may differ due to the influence of soil and vegetation on temperature and water availability.

We tested the overarching hypothesis that NRM is an important, year-round source of moisture in xeric and mesic grasslands by 1) offering a first-time quantification of NRM’s contribution to annual mass loss, relative to rain, 2) describing the factors that control NRM decomposition, and the conditions under which it occurs and 3) assessing the ability of different approaches to estimate NRM frequency and NRM decomposition.

We took a coupled empirical-modeling approach to meet these goals. We first quantified NRM type, frequency, and duration, and measured microbial respiration and mass loss of surface litter under NRM at three grassland locations with contrasting moisture regimes (a hyper-arid site in the western Namib Desert with high NRM; an arid site in the eastern Namib Desert with very infrequent NRM; and a mesic site in an Iowan grassland which experiences both high rainfall and NRM). These empirical measurements allowed us to assess the conditions under which NRM decomposition occurs, and develop predictive relationships between NRM meteorology and decomposition. Using this information, we modeled annual C loss when excluding and including NRM (in addition to rain) at each site. We applied our model that predicted annual C loss attributed to rain and NRM to an additional site, Sevilleta, New Mexico, to test the robustness of our estimate of NRM decomposition at a site with intermediate rainfall (semi-arid grassland).

## Methods

### Site descriptions

Our entire study (NRM characterization, CO_2_ flux measurements, and modeling) included analysis efforts in three regions: the Namib Desert (Namibia), Iowa tallgrass prairie (USA) (**Fig. 1**, **Table 1**), and a New Mexico semiarid grassland (USA) (**Fig. S1**, **Table 1)**. We took empirical measurements (litter flux and mass loss, and direct measurement of NRM) at two sites in the Namib with contrasting moisture regimes, and one site in Iowa. We chose sites in the Namib (hyperarid desert) because we have ongoing investigations of microbially-mediated surface litter decomposition here that are facilitated by existing meteorological infrastructure monitoring NRM. The mesic grassland site in Iowa (tall-grass prairie) was chosen because it provided an extreme contrast to the Namib sites, and because of its close proximity to one of our home institutions. We also analyzed data from a semiarid site, Sevilleta, New Mexico to assess whether NRM is likely to be important in regions with rainfall intermediate to the Namib and Iowa, and to test approaches for characterizing NRM decomposition using long-term meteorological records which lack leaf wetness sensor data.

**Figure 1.**
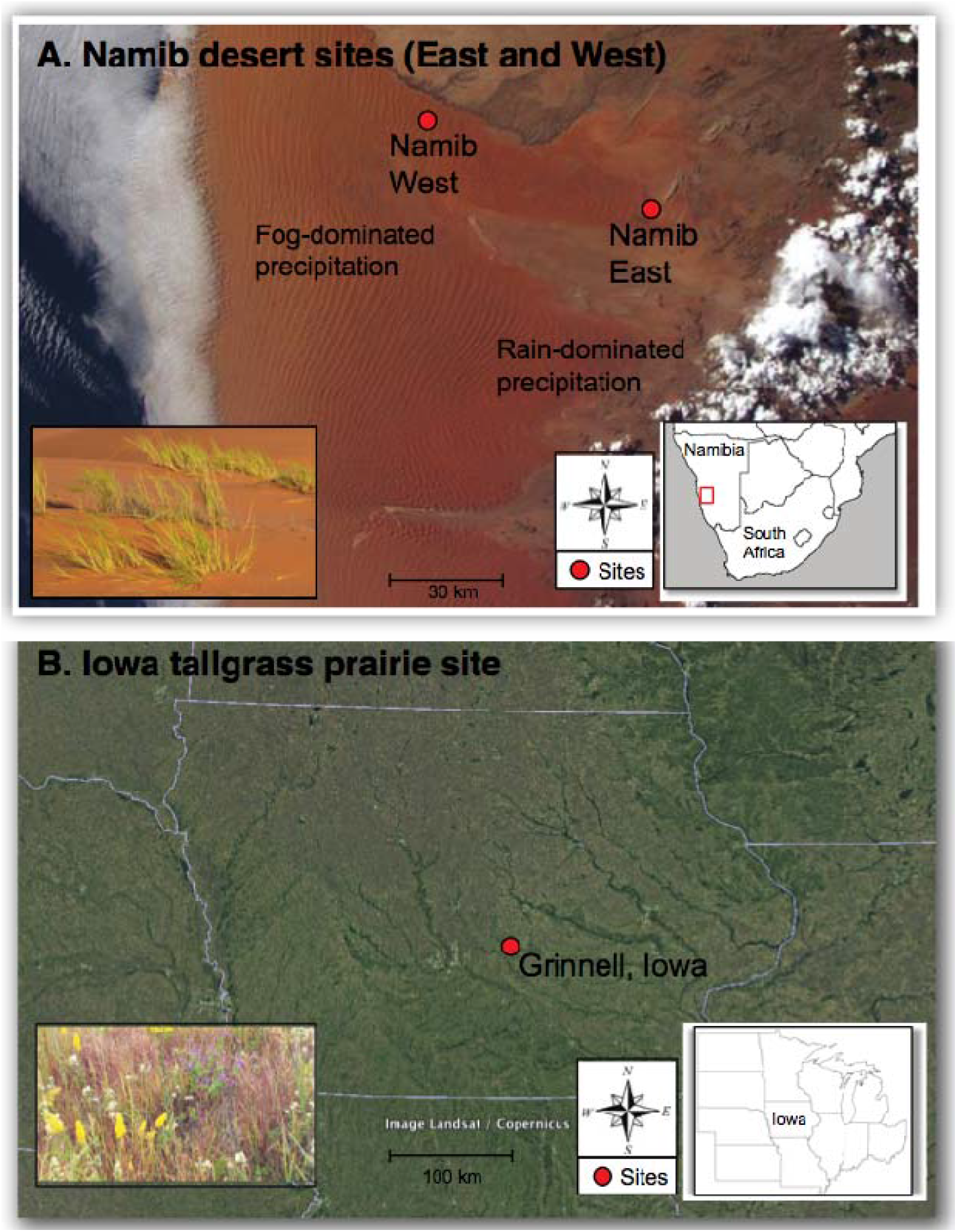
Site descriptions. This study was conducted in: A: the Namib Desert, Namibia, at the NRM-dominated ‘Namib West’ site and rain-dominated ‘Namib East’ site. We measured litter flux from *Stipagrastis sabulicola*, the dominant plant (inset). B: Iowa tallgrass prairie in Grinnell, Iowa. We measured litter flux from *Andropogon gerardii*, the dominant plant. Inset shows diverse mix characteristic of tallgrass prairie.

**Table 1.** Meteorological and site details for 4 sites studied. We measured NRM frequency, respiration, and mass loss at Namib East, Namib West, and Iowa, and measured NRM frequency at Sevilleta, modeling mass loss.

|  | Namib East | Namib West | Iowa | Sevilleta |
| --- | --- | --- | --- | --- |
| Site coordinates | S 23.7835<br>E 15.7796 | S 23.5604<br>E 15.0410 | N 41.7568<br>W 92.7151 | N 34.3592<br>W 106.691 |
| Met record dates | 6/24/15 - 6/4/16 | 6/15/15 - 6/16/16 | 3/9/16 - 1/6/17 | 1/1/11 - 1/1/16 |
| Met record length (d) | 346 | 367 | 303 | 1825 |
| Mean annual temp (°C) | 23.1 | 21.0 | 13.1 | 13.8 |
| Mean annual rainfall (mm) | 81 | 19 | 971 | 240 |
| Mean RH (%) | 32 | 49 | 69* | 40 |

<sup>±</sup>https://www.currentresults.com/Weather/Iowa/humidity-annual.php

The Namib sites are located in a linear dune system, and include an east and west site that differ in rain and fog inputs (**Fig. 1A**). At the Namib East site, rainfall is more prevalent (~81 mm), and fog is rare (Lancaster and others 1984; Eckardt and others 2013). Dew frequency had not been quantified at the east site before this study. At the Namib West site, mean annual rainfall is low (~25 mm), and fog and dew are common (each occurring >50 nights per year) (Henschel and Seely 2008; Eckardt and others 2013; Jacobson and others 2015). Both Namib sites are dominated by the dominant perennial dunegrass *Stipagrostis sabulicola* (**Fig. 1A inset**). The Iowa site is in a restored tallgrass prairie near Grinnell, Iowa, USA with a mean annual rainfall of 971 mm (City of Grinnell) (**Fig. 1B**). NRM frequency had not been quantified before this study. Vegetation is dominated by *Andropogon gerardii* (**Fig. 1B inset**) and a diverse assemblage of prairie forbs. The New Mexico site is a semiarid grassland in the Sevilleta National Wildlife Refuge with a mean annual rainfall of 240 mm (Peters and Yao 2012) (**Fig. S1**, **Table 1**). NRM frequency had not been quantified before this study. Notably, at this site Vanderbilt et al. (2008) found that mass loss correlated poorly to precipitation in their long-term study, suggesting alternative decomposition mechanisms are at play. We made no empirical measurements at the site, but analyzed NRM frequency from standard meteorological data (http://digitalrepository.unm.edu/lter_sev_data/8/). Vegetation here is dominated by *Bouteloua eriopoda* and *B. gracilis*.

### Meteorological measurements and analysis of NRM frequency using leaf wetness sensors

We assessed meteorological conditions at Namib West, Namib East, Iowa, and Sevilleta sites (**Table 1**) taking advantage of existing infrastructure and datasets, and adding capabilities where necessary. Namib West is equipped with a SASSCAL meteorological station (http://www.sasscal.org/), which houses a Campbell CS215-L temperature and humidity probe positioned at 2 m, a Juvik fog collector, and a leaf wetness sensor (Campbell 237-L, uncoated, 10 cm height). The Juvik fog collector is an omnidirectional, cylindrical aluminum fog screen, positioned at a height of 1.5 m (Juvik and Nullet 1995), and fitted onto a Young tipping rain gauge (Y52203, Young Company, Michigan, USA). At Namib East, we monitored air temperature and relative humidity (RH) (height: 150 cm), rainfall and leaf wetness (height: 25 cm) using a HOBO data logger and sensors (H21-002, S-THB-M002, Davis S-RGD-M002, S-LWA-M003) (Onset Computer Corp., USA).

A similar meteorological array was used at the Iowa site, except that the RH and temperature sensors were positioned at a height of 65 cm, in the midst of the prairie vegetation. An automated tipping-bucket rain gauge (HOBO, RG3-M, 15.24 cm diameter, 0.2 mm resolution) was placed nearby at an elevation of 1.5 m. In Sevilleta, New Mexico, we analyzed NRM frequency from data recorded at the Deep Well Meteorological Station (No. 40), including hourly RH, rainfall, and air temperature. RH and temperature sensors were positioned at a height of 2.5 m. Instrumentation details can be found at http://digitalrepository.unm.edu/lter_sev_data/8/. Leaf wetness data were not recorded at Sevilleta.

We estimated total wet hours at a site by using either (1) the number of hours leaf sensors were wet, (2) the number of hours that exceeded an RH threshold (Sentelhas and others 2008) and (3) a function (‘likelihood wet’) that estimated the likelihood a sensor would be wet, based on RH. We attributed a wet hour to rain if rainfall was detected during that hour.

Leaf wetness sensors (**Fig. 2E**) have been widely used by plant pathologists to estimate periods of wetness that are independent of rainfall (Rowlandson and others 2015), and in other studies to estimate NRM (Gotsch and others 2014; Gliksman and others 2017) by measuring water droplets and films on electronic grid surfaces. In sensors that measured wetness on a discrete scale (Campbell 237-L, Namib West), the wet-dry transition occurred at ~150 kohm; for continuous-scale (%) wetness sensors (Iowa, Namib East), we conservatively defined ‘dry’ periods as those below 10% wetness. We estimated wet hours from RH by totaling hours that RH exceeded either 75% RH (low threshold) or 90% RH (higher threshold), as informed by previous work (Sentelhas and others 2008). Finally, we determined the relationship between wetness sensor readings and RH at each site, developing a function (“likelihood wet”) for the likelihood that the leaf wetness sensor indicated ‘wet’ for a given RH. These likelihood curves were remarkably similar across sites (**Fig. S3**), justifying use of the mean curve to estimate the number of hours in each site that the sensor was wet (with an uncertainty band based on the between-site variation), including the Sevilleta site, as derived from RH. This estimate of wet hours was used to calculate C loss (see last section).

**Figure 2:**
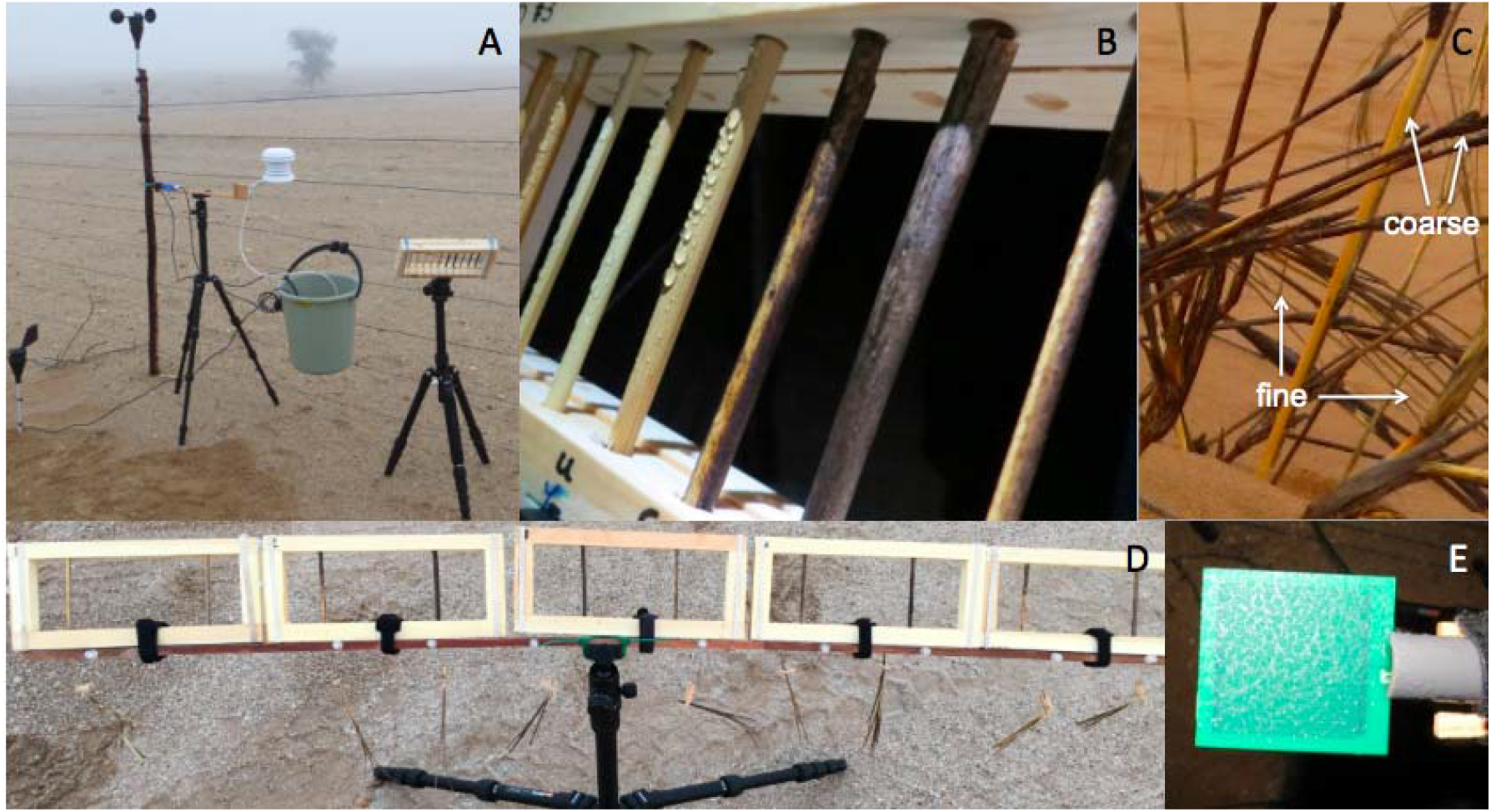
Photographs of standing surface litter and methodologies. A: Measurement of litter gravimetric moisture and flux in litter rack, and associated portable meteorological station at Namib West. B: Litter rack mimicking standing coarse litter *in situ*, shown with droplets from NRM. C: Different litter types: fine (leaves) and coarse (tillers, >2mm diameter), shown here on *S. sabulicola*. D: Fine litter hanging below coarse litter racks during NRM exposure. E: Leaf wetness sensor containing condensed water during an NRM event.

Having defined wet hours (as determined by leaf wetness in Iowa, Namib and likelihood function in Sevilleta), we determined the mean temperature and humidity associated with NRM and Rain within each site during NRM events. We were also interested in the duration of a typical rain and NRM event, which required that we delineate the start and end to an event. In our analysis, events began by at least 2 wet hours in a row (to exclude spurious Wet hours) and ended at the first 2 dry hours detected. Delineation of events was not possible in Iowa because leaf wetness sensors were often continuously wet for weeks at a time (likely due to the location of the LWS within the prairie canopy). Thus, in contrast to the drier sites, many ‘events’ at the Iowa site could have included both rain and NRM. Please see https://github.com/ktoddbrown/NRM_litter_decay for the code associated with this analysis.

### Empirical measurements of mass loss and respiration in the field

We measured litter mass loss using litter racks (**Fig. 2B)** instead of traditional litter bags, which we found can alter NRM (See Supplement for full details justifying this method). We deployed pre-weighed native coarse tillers (4-6 x 90 mm) in these wooden racks at the Namib and Iowa sites at ~0.5 m height. In the Namib sites, we monitored mass loss of *S. sabulicola* standing litter that was collected after senescence from each site, air-dried, and stored at room temperature until rack deployment. In Iowa, *A. gerardii* was collected in the fall following senescence, dried at 35°C, and stored at room temperature until rack deployment. After a one-year deployment in racks mounted on poles at each site, tillers were similarly dried and stored individually in air-tight Whirlpack bags until weighed. Mean percent mass loss of the tillers (n=4-10) was compared across sites using a 1-way ANOVA.

In addition to mass loss, we assessed flux rates and moisture content of litter under NRM events. We examined ‘coarse’ (thick tillers, ~5 mm diameter, used in mass loss studies) as well as ‘fine’ (stem sheaths and leaves) litter types (**Fig. 2C**) to test whether the effect of NRM differed by substrate (Jacobson and others 2015). Tillers were collected for respiration measurements in the same way they were collected for assessing mass loss (see above). We deployed racks on a tripod in the evening hours, after dark, when climatic conditions suggested an NRM event might occur (**Fig. 2A,C**). We also deployed an autoclaved subset of coarse litter ‘controls’ to test whether the majority of respiration was microbially-mediated, or possibly mediated by abiotic mechanisms such as photolysis or thermal emission after sunrise (Lee and others 2012; Day and others 2019). Tillers were kept sterile and in the dark until deployed, but some respiration could still be microbial in origin since we could not prevent sterile tillers from being colonized by airborne inoculum during an NRM event (Evans and others 2019). Fine litter (<1 mm x 4-10 mm x 80-120 mm, **Fig. 2C**) was suspended by small clips from a litter stand directly below the racks when an NRM event was anticipated (**Fig. 2D**).

At each measurement time point, we first extracted and weighed individual litter pieces to determine gravimetric moisture content. Then flux from each tiller was measured over a 3-minute period (including a 30-s dead band period) using a Li-8100 CO_2_ Flux system (LI-COR Inc., Lincoln, NE), equipped with a small (~55 ml) respiration chamber (LI-COR 6400-89). The majority of flux measurements were made at night when it was dark and cool (or, after sunrise, at temperatures <25°C and out of direct sunlight) so photolysis and thermal degradation was unlikely or minimal. After measurement, litter pieces were then immediately replaced in the rack or stand. At the conclusion of the NRM event, litter was dried (at 35°C) to determine gravimetric moisture, and flux was expressed on a dry weight basis.

We first analyzed whether respiration observed under NRM was microbial in origin by comparing flux rates of sterilized to unsterilized pieces of litter (t-test, n=5-10). We tested controls for NRM respiration and gravimetric moisture using multiple linear regression. We included all replicate litter pieces in a sampling time point after finding no significant effect of rack (p>0.1) or event (p>0.1), and excluding points at the end of events, which were under-sampled (see Results). With this dataset (N=128), we tested (1) the effect of site, gravimetric moisture, and litter type, on respiration; and (2) the effect of leaf wetness and litter type on gravimetric moisture at Iowa and Namib sites. Since C flux at Namib East and West sites did not differ in response to any of these environmental drivers, we combined into one ‘Namib’ site. All statistical analyses were performed in R v. 3.4.0 (R Core Team 2017).

### Extrapolation of C loss across space and time

We assumed that microbially-mediated decomposition occurred during wet periods at all sites, as supported by our field observations. We used our empirical measurements of gravimetric moisture and litter respiration to determine the C flux associated with a wet hour. We calculated the mean C flux (with 95% confidence interval) when litter was above 15% gravimetric moisture (an approximate threshold for respiration turning ‘on’ across sites, see **Fig. 4A**), and estimated C loss at all sites by multiplying this flux rate by estimated wet hours (see above for approaches for quantifying wet hours). We were unable to directly correct for temperature in our study (e.g. using a Q10 sensitivity), and suggest future studies do so. However, we measured flux under a relatively broad temperature range, and the uncertainty propagates in confidence intervals for mass loss estimates. To facilitate comparisons across sites, which had slightly different measurement periods, we converted estimates to an annual scale. To enable qualitative comparisons to measured mass loss, we converted C to litter mass by assuming 50% of mass was C (Schlesinger 1977).

## Results

### Characterization of non-rainfall moisture across sites

Despite the large difference in rainfall across the sites (**Table 1**), many aspects of NRM were similar. For instance, the proportion of wet hours attributed to NRM was exceedingly high (85.0-99.1%), and NRM generally occurred during humid (81%-93%) and cool (12-13°C) periods for several hours or more (**Table S3**, **Fig. S2**), conditions sufficient to induce microbial activity. We observed substantially more total NRM wet hours compared to rainfall-wet hours at all sites. In the Namib sites, temperature during NRM was generally lower than it was during rain, and RH was higher (**Table 2**, **Table S2**). In Iowa, NRM occurred across a broader range of temperatures than in the Namib (**Table 2**), and at more similar temperatures to those in rain periods. In addition to their far greater frequency, NRM events may also last longer than rain events (**Fig. S2**), but we could not test this comprehensively because of the few rain events in the Namib sites, and the challenge in delineating events in the Iowa site. (Specifically, wetness sensors measured long periods (multiple day) of wet hours in Iowa, especially in the summer months, because of the consistently high humidity at the height of the sensor (65 cm) resulting from the dense vegetation canopy that traps soil-derived moisture.) In the Namib, NRM events were longer in Namib West (7.3 h) than Namib East (6.0 h) (p=0.007) (**Fig. S2**). A general caveat to these trends is that our sampling period was a single year, not long-term mean annual NRM frequency. We have no reason to assume our NRM data are unique to this year and note that annual precipitation means for our sampling periods are similar to or slightly lower than published long-term means at each site (**Table 1**).

**Table 2.**
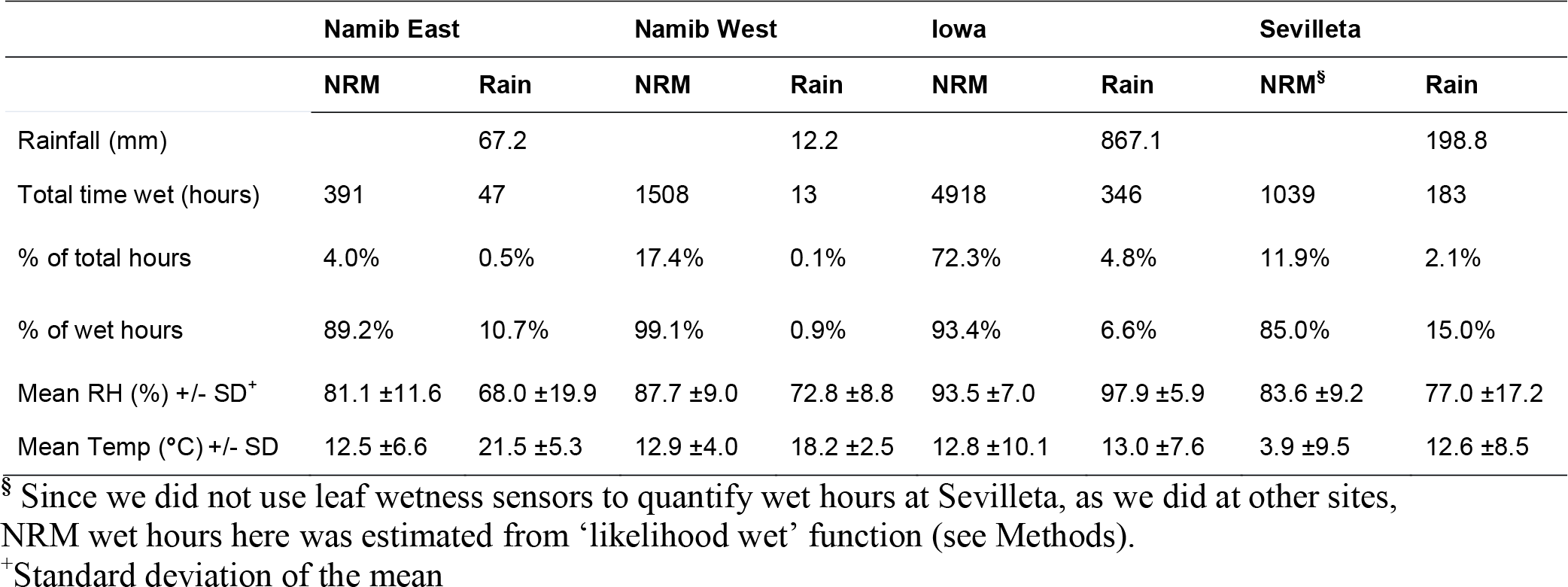
Summary of non-rainfall moisture (NRM) and rain across sites. Wet hour indicates an hour when a leaf wetness sensor is wet (see Methods for approach at Sevilleta), either due to NRM (left) or rain (right). Data are standardized by dataset length and reported on a per year basis to facilitate comparisons across sites.

|  | Namib East |  | Namib West |  | Iowa |  | Sevilleta |  |
| --- | --- | --- | --- | --- | --- | --- | --- | --- |
|  | NRM | Rain | NRM | Rain | NRM | Rain | NRM<sup>§</sup> | Rain |
| Rainfall (mm) |  | 67.2 |  | 12.2 |  | 867.1 |  | 198.8 |
| Total time wet (hours) | 391 | 47 | 1508 | 13 | 4918 | 346 | 1039 | 183 |
| % of total hours | 4.0% | 0.5% | 17.4% | 0.1% | 72.3% | 4.8% | 11.9% | 2.1% |
| % of wet hours | 89.2% | 10.7% | 99.1% | 0.9% | 93.4% | 6.6% | 85.0% | 15.0% |
| Mean RH (%) +/- SD<sup>+</sup> | 81.1 ±11.6 | 68.0 ±19.9 | 87.7 ±9.0 | 72.8 ±8.8 | 93.5 ±7.0 | 97.9 ±5.9 | 83.6 ±9.2 | 77.0 ±17.2 |
| Mean Temp (°C) +/- SD | 12.5 ±6.6 | 21.5 ±5.3 | 12.9 ±4.0 | 18.2 ±2.5 | 12.8 ±10.1 | 13.0 ±7.6 | 3.9 ±9.5 | 12.6 ±8.5 |

<sup>§</sup> Since we did not use leaf wetness sensors to quantify wet hours at Sevilleta, as we did at other sites, NRM wet hours here was estimated from ‘likelihood wet’ function (see Methods).
<sup>+</sup>Standard deviation of the mean

The different approaches for estimating wet hours (wetness sensor, high humidity, and a likelihood function) were generally comparable within a site, and consistently estimated more wet hours due to NRM than wet hours attributed to rain (**Fig. 3**). Estimates of wet hours from leaf wetness sensors fell within the range of estimates generated using RH threshold, but the RH threshold chosen (75% vs. 90%) had a large impact on the proportional contribution of NRM to wet hours in a site (**Fig. 3**). RH of 85%, which is supported by measured thresholds for fungal activity (Dix and Webster 1995), produced estimates near those measured by leaf wetness sensors. A “likelihood wet” function also produced wet estimates similar to those measured by leaf wetness at each site (**Fig. S3**, **Fig. 3**), which also indicated that our estimates of wetness frequency at Sevilleta were similar to what we would have measured with a leaf wetness sensor.

**Figure 3.**
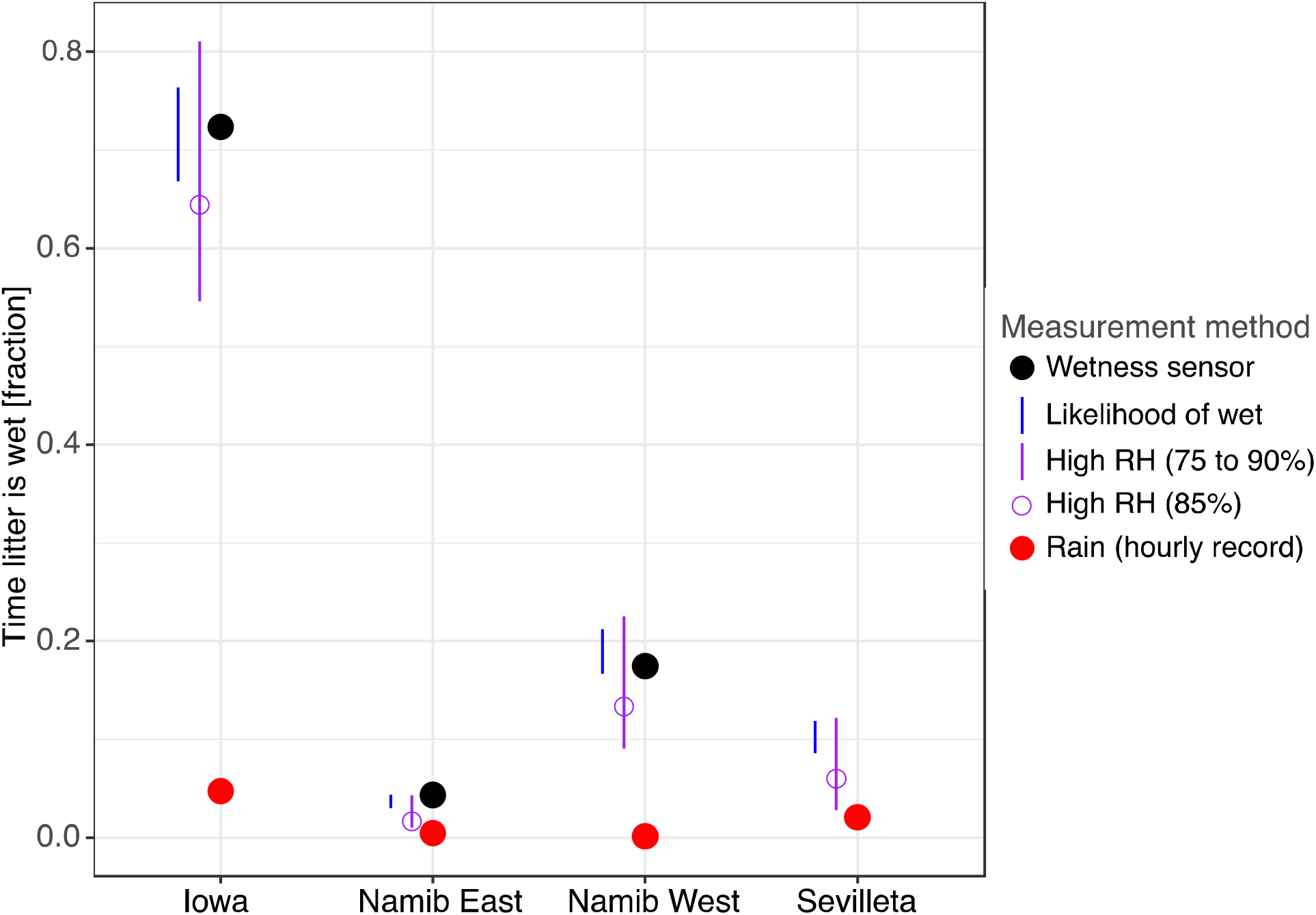
Fraction of time litter is wet, estimated by different techniques, and used to extrapolate mass loss. Red dot: raining time. Black dot: wet hours as estimated by leaf wetness sensors. Blue line: likelihood of a wet sensor (‘likelihood wet’ function) for a given relative humidity, based on relationships at Iowa and the Namib. Purple line: estimates using RH threshold, with the lower bound using a threshold of 75% and upper bound, 90%, and open purple circle showing 85%.

**Figure 4:**
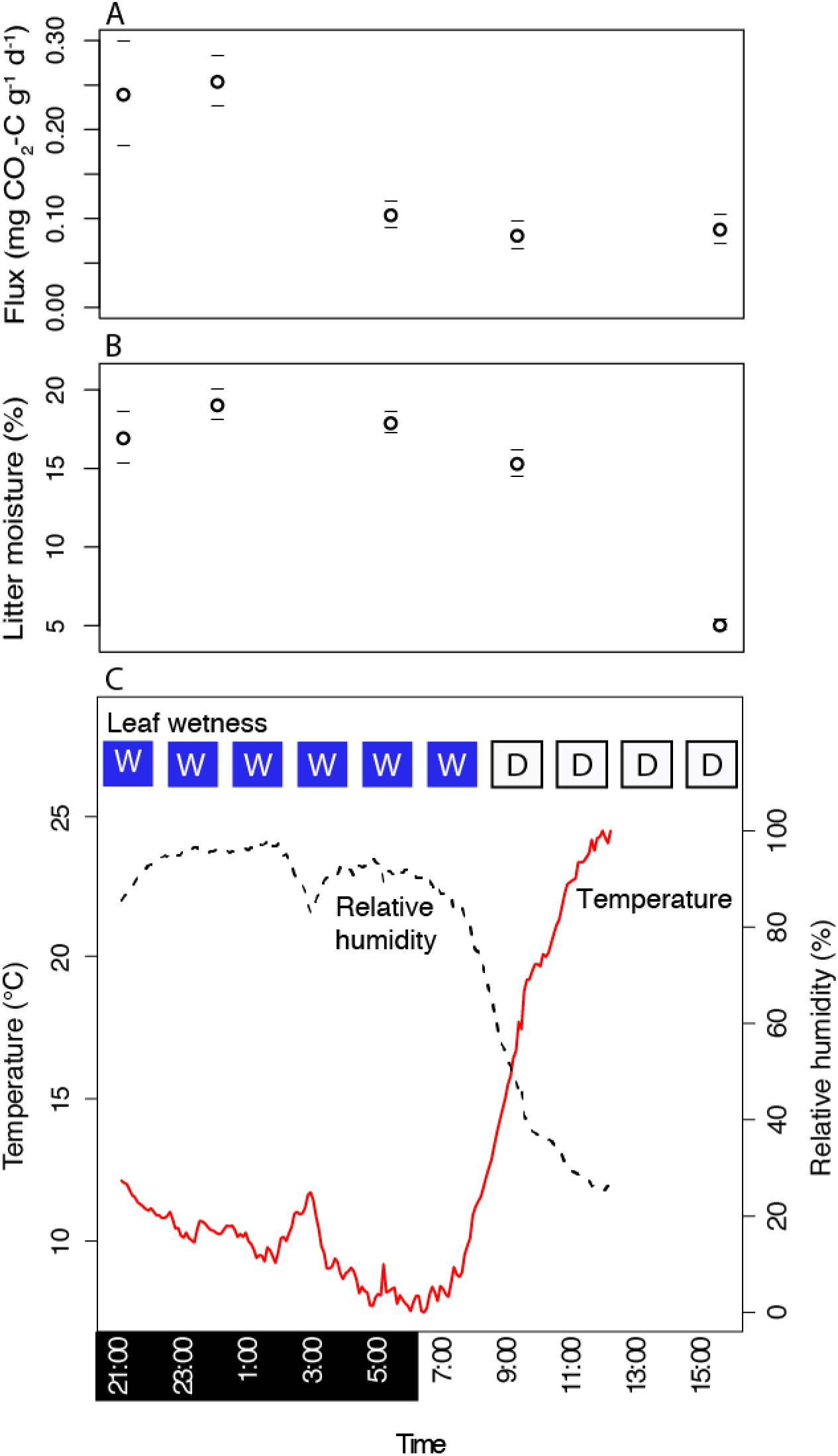
Response of standing *S. sabulicola* (coarse) litter to one dew event at Namib West on 3 June 2015 (see Table S3 for all events)). A: litter respiration, B: gravimetric moisture (n=10, dashes represent 1 SE above and below the mean represented by symbols), and C: meteorological parameters over the course of one night (W=wet and D=dry leaf wetness reading). Respiration shown for coarse litter; fine litter had higher rates of respiration (1.81 mg CO_2_-C/g litter/day at 00:45, Table S3). Dew began around 19:00, when leaf wetness read “slightly wet” and relative humidity was 83%.

### Field measurements of NRM-induced litter respiration

We observed significant CO_2_ flux under multiple NRM events from standing litter in both arid and mesic grassland systems (**Table S3**). In a typical NRM event in the Namib that induced respiration (**Fig. 4**), flux began during the night as temperatures decreased and relative humidity (RH) increased. Flux was sustained with high litter moisture during the night-time hours, then decreased in the morning as RH decreased and temperature increased (**Fig. 4B & C)**. Flux rates at a single time point were as high as 2.63 mg CO_2_-C/g litter/day (mean across N=5 pieces of fine litter during fog in Namib West) (**Table S3**). The majority of CO_2_ flux was mediated by microbial activity; sterile tillers exposed to NRM had very low flux that was significantly lower than respiration from nonsterile tillers (**Table S4**). Since it was difficult to predict when dew would occur, we started most flux measurements in the middle of an event (**Fig 4A**), so we know less about moisture levels that induce respiration under NRM. (We do, however, have evidence from the lab that this occurs very rapidly when gravimetric moisture surpasses a 10-15% threshold (Jacobson and others 2015).) Although events generally ended by mid-morning (09:00) (**Fig. 4C**), on three occasions we observed tillers that were slightly wet (5-10% gravimetric moisture) and respiring at low levels into the late morning and early afternoon, even though the leaf wetness sensor measured zero (**Table S3**, **Fig. 4**). Thus, extrapolations based on leaf wetness alone could underestimate NRM decomposition.

We used a regression approach to test the generality of the response of respiration to NRM across litter type, site, and precipitation type (rainfall, fog, dew, high humidity) in the Namib and Iowa. Since we were interested in controls on the maximum and sustained respiration flux, we excluded all flux measurements that occurred while litter was drying (e.g. at the end of an event) from regression analysis (**Table S3**, right column). NRM induced significant respiration at Namib West (where fog is common), but also at Iowa and Namib East sites (**Fig. 4**, **Table S3**), verifying that microbial activity under NRM is not unique to sites where fog is frequent, or in arid systems alone.

Gravimetric moisture explained 60% of respiration under NRM across sites (p<0.001) (**Fig. 5A)**, although explained little variation in Iowa (y=0.005x+0.669, R^2^=0.06, p<0.04), compared to the Namib (y=0.02x+0.17, R^2^=0.7, p<0.001, **Fig. 5A**). There was no difference in flux response between the two Namib sites. The slope of respiration response differed between Namib and Iowa sites, however (p<0.05 to reject the null of equal slopes). Fine litter flux in Iowa was more constrained at the wetter end, but this may be explained by the fact that sampling in Iowa took place during cooler events (mean temperature for fine litter NRM events in Iowa=8.4°C and Namib=14.9°C, **Table S3**), rather than differences in microbial community activity across sites. Litter type significantly affected the extent of litter wet up during NRM (p<0.001). Fine litter (leaves and tiller sheaths, see **Fig. 2C**) became wetter than coarse litter (tillers) under the same leaf wetness (**Fig. 5B**), and on average achieved higher flux (**Fig. 5A**). Greater flux in fine litter was also due to its lower moisture threshold for initiating respiration (5% vs. 13%, respectively, p<0.05 to reject null of equal y-intercept).

**Figure 5.**
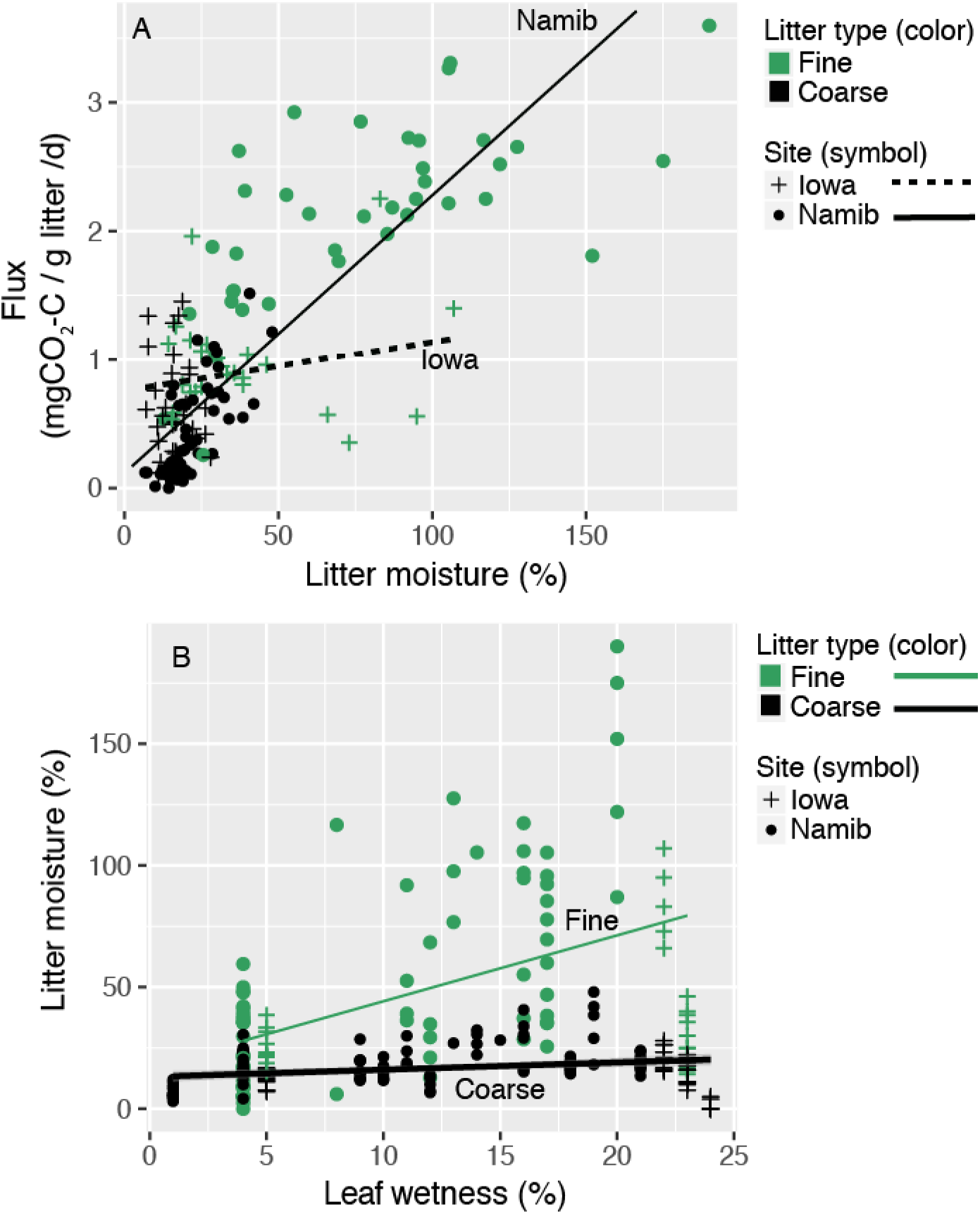
(A) Under NRM, gravimetric moisture was positively related to respiration for the Namib (combined East and West, y=0.02x+0.17, R^2^=0.7, p<0.001) and Iowa (y=0.005x+0.669, R^2^=0.06, p<0.04). (B) Under NRM, gravimetric moisture content of fine litter increased significantly more than that of coarse litter under the same meteorological conditions (here shown by leaf wetness sensor) (reject null of equal slope, p=0.01).

We did not measure flux under rain in enough rain events to assess this statistically, but our data suggest that NRM events result in at least as much wet-up and C loss as rain events. During the rain event we documented in Iowa, mean flux was 0.68 CO_2_-C/g litter/day (N=5), within the range of flux observed under NRM events (0.004 – 0.91 CO_2_-C/g litter/day, **Table S3**). During a relatively large rain event at the Namib West site (12.8 mm, 6 June 2016), coarse litter gravimetric moisture was similar to moisture reached under typical NRM events (maximum 32% under rain, 35% under NRM), although fine litter did not get nearly as wet under rain (maximum 20.5% under rain and 145% under NRM) (**Table S3**). We did not discern any differences in moisture or flux patterns between NRM types (fog and dew; p-value > 0.05, N=5 dew and N=3 fog events).

### Contribution of NRM to annual decomposition

Litter mass loss, measured empirically, was highest in Iowa, and generally low in the arid and hyper-arid Namib sites (**Fig. 6**). Notably, mass loss in the in Namib West was similar to (and even trending higher than) mass loss in Namib East (p=0.66), a site with more rainfall, but less NRM (**Table 2**, **Fig. S2**). The exclusion of NRM (that is, using rain as the only driver of decomposition) resulted in very low estimates of annual mass loss at all sites (**Fig. 7)**. Incorporating NRM resulted in a ~6-fold increase in estimated C flux at the most mesic Iowa site, to a >100-fold increase at the hyper-arid Namib West site (**Fig. 7**). The height of the sensors in Iowa (which were beneath the plant canopy, unlike sensors at other sites) may have contributed to the high NRM measured at this site. Using rainfall hours alone underestimated observed mass loss in the sites where it was measured (Namib and Iowa, lines, **Fig. 7**, standardized to annual scale), while NRM+Rain extrapolated estimates fell within the range of observed values. This is true even though extrapolation efforts did not technically include photodegradation (photolysis or photopriming); modeling was based on respiration rates made on litter stored in the dark and deployed at night. This could have contributed to underestimation of observed mass loss at high-UV sites like Namib East. At Sevilleta, when NRM was included, mass loss estimates were definitely closer to observed values (10% for *A. gerardii* (Brandt and others 2010), 20.1% for *B. eriopoda*, (Vanderbilt and others 2008)), but in these studies, litterbags were used in this study which could underestimate observed NRM decomposition (see Supplemental Methods).

**Figure 6.**
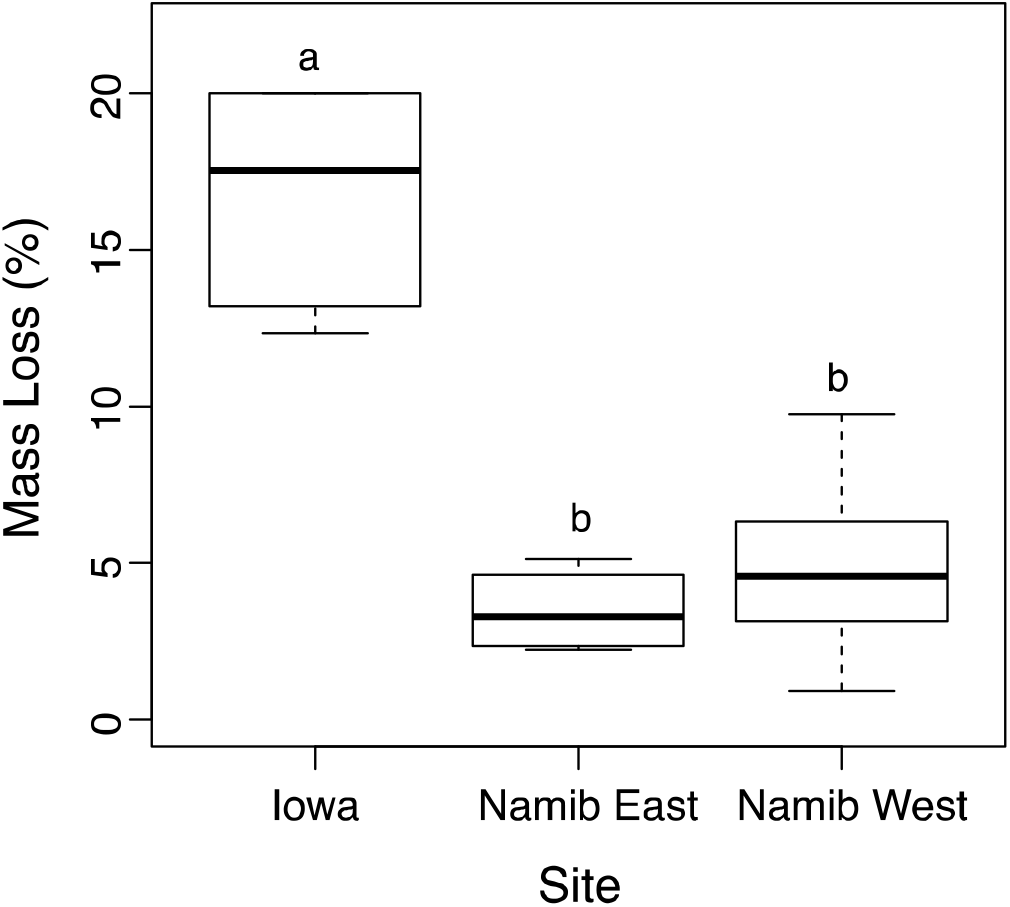
Mass loss of standing litter in mesic (Iowa) and hyperarid (Namib) sites that had different rain and NRM regimes. Box shows upper and lower quartiles and line within the box represents the median. Litter was native (*S. sabulicola* in Namib sites, *A. gerardii* in Iowa) coarse grass ‘tillers’ deployed in standing litter racks at the height of native standing litter. Different letters represent significant (p<0.01) differences (pairwise t-tests) among mean mass loss in Iowa (N=5, 303 days deployment), Namib East (N=5, 343 days), and Namib West (N=26, 344 days).

**Figure 7.**
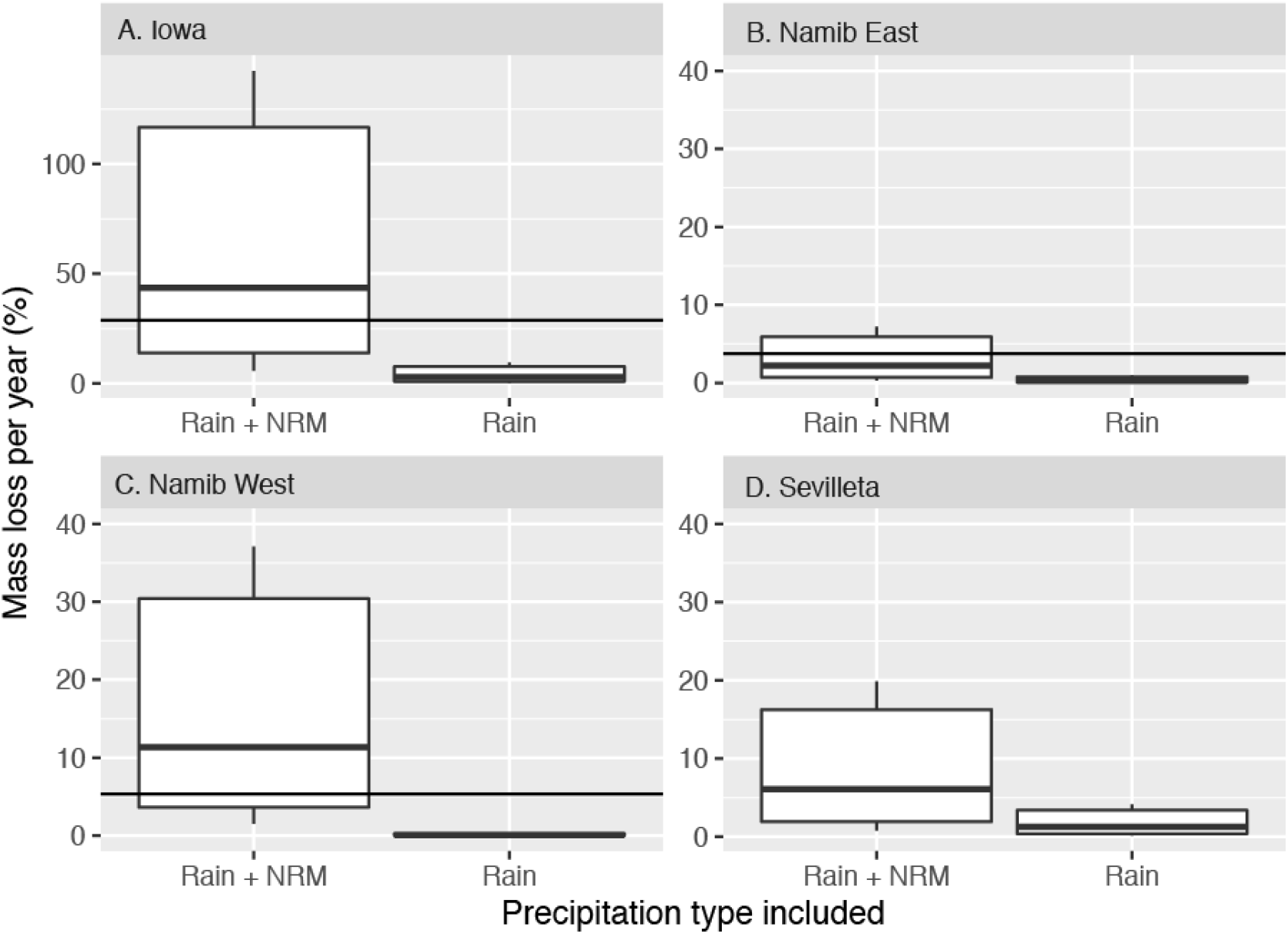
Estimated litter mass loss when NRM-decomposition is excluded (“Rain”) and included (“Rain+NRM”). Black solid lines show observed mean mass loss at each site, standardized to an annual scale to facilitate comparison. (Note Fig. 6 values show the median, not standardized to annual)

Mass loss estimates that included NRM had large confidence intervals (**Fig. 7**). The primary source of this uncertainty was the wide range of potential flux rates that can be induced under wet conditions (refer to data in **Fig. 5**), rather than uncertainty surrounding estimation or extrapolation of NRM frequency. Even when NRM frequency was directly measured using leaf wetness sensor, confidence intervals were large for the overall carbon loss rates (**Fig. S4C).** Still, other factors did introduce some variation in estimates. Specifically, there were some differences in the relationship between leaf wetness and RH at different sites; for example sensors became wet at slightly lower RH values at Namib East than at Namib West (**Fig. S3**). However, this across-site variation in wet hour estimates contributed little error to mass loss estimates compared to the respiration uncertainty (**Fig. S4B**). The global function (**Fig. S3**) was also in line with previous estimates (e.g. predicted sensors were more likely to be wet than dry around 82% RH (Sentelhas and others 2008)). Finally, as noted above, we also examined the accuracy of estimating wet days using an RH threshold approach. While we did not use this approach for our primary extrapolation of mass loss (in **Fig. 7**), we did find that the RH threshold chosen is extremely important. Decomposition estimates were very sensitive to the threshold value chosen (75%-90% in this study, **Fig 3 and Fig. S4**), again highlighting the need for site-specific calibrations of wetness sensor-RH relationships.

## Discussion

### NRM contributes to annual mass loss of standing litter across grassland types

Our empirical measurements demonstrated that NRM (fog, dew, high humidity) is an important, year-round driver of standing surface litter decomposition in sites representing the extreme ends of grassland systems (hyper-arid and mesic), and that similar NRM events that induce microbial activity are frequent in semi-arid grasslands as well. We estimated that in all sites, 85-99% of wet hours were attributable to NRM, and as a result (and as informed by our on-the-ground measurements of respiration under these periods), NRM was a large contributor to annual decomposition of surface litter at all sites – greater than that of rainfall. Although our goal was not to produce a predictive decomposition model – this will require larger empirical datasets, and incorporation of other factors like temperature – our first effort to scale contributions of NRM does show that including NRM produced values much closer to measured mass loss. Overall, our extrapolation demonstrates what many other studies have suggested (Dirks and others 2010; Jacobson and others 2015; McHugh and others 2015; Gliksman and others 2017): that NRM is not just a fleeting stimulator of occasional respiration, but rather an important driver of surface litter decomposition on an annual scale, in many grassland types.

In dryland sites, many decomposition models that use rainfall as the sole moisture source underestimate empirical observations of surface litter mass loss (Parton and others 2007; Adair and others 2008; Brandt and others 2010; Currie and others 2010), even though it is an important predictor of mass loss in more mesic systems. Our study suggests that exclusion of NRM from models could contribute to this underestimation. This is highlighted by our empirical measurements: one year of mass loss in a site with almost no rain but high NRM (Namib West) had slightly higher mass loss than another site with higher rainfall but lower NRM (Namib East). Furthermore, mass loss estimates were substantially closer to observed values when NRM was included in our model extrapolation. Other mechanisms, in particular photodegradation, are also likely to be important in dryland decomposition, and have improved model predictions of dryland decomposition (Brandt and others 2010; Adair and others 2017). Photodegradation may be an especially important stimulator of decomposition when it interacts with – and facilitates – microbial decomposition (Foereid and others 2010; Gliksman and others 2017; Day and others 2018); in fact, the contribution of high-UV periods to decomposition may be negligible without intermittent, microbially-active wet periods (Lin and others 2018), at least as long as the system is generally moisture-limited (Smith and others 2010). Our study shows that NRM could provide these wet periods that induce microbial activity, as suggested by Jacobson et al. (2015) and Gliksman et al. (2016). We found that NRM delivers these essential wet periods on a diel scale, and contributes more wet hours for microbial activity than rainfall, which may not be the best indicator of water availability.

NRM was also the primary contributor to wet periods in our mesic grassland site (93% of total wet hours), highlighting the ubiquity of NRM-induced wetness across grassland systems. We found that excluding these periods in our rain-only model resulted in mass loss estimates much *lower* than observed values, which is seemingly at odds with the relatively good performance of traditional (rain-driven) decomposition models in mesic grasslands (Parton and others 2007; Adair and others 2008). We suspect that this is because relative humidity (RH) is included in many traditional models, thus implicitly allowing NRM to influence water availability in soils and litter (e.g. Parton et al. (2001)); whereas our rain-only extrapolation did not. An implicit approach might be sufficient to predict RH-induced wetness that is due to retention of moisture (through reduced evapotranspiration) in the soil-grass canopy system. However, this approach would not capture NRM decomposition resulting from shorter-term (e.g. diel) RH fluctuations, which are frequent in xeric systems.

### Controls on NRM decomposition of surface litter

Our empirical measurements of NRM-induced respiration in the field show that moisture thresholds under NRM are similar to those observed in previous studies and in the laboratory. Respiration ‘turned on’ under NRM around 13-20% gravimetric moisture (depending on litter type), which narrows our previous estimates (10.5-30%), and is remarkably close to minimum thresholds for initiation of litter respiration reported in previous laboratory studies (10-20%) (Bartholomew and Norman 1947; Nagy and Macauley 1982) and under high humidity in the field (10%) (Gliksman and others 2017). Thresholds for initiating vs. ceasing respiration may differ due to physical properties of the litter (e.g. coarse tillers vs fine litter), physiological controls on microbial community resuscitation and desiccation, or how litter wets and dries relative to the distribution of microbial biomass, which changes as litter ages (unpublished data, Logan et al. in prep).

Our findings reiterate that NRM frequently induces moisture levels sufficient for microbial activity, and standing litter will respire when sufficiently moist, no matter if from rain or NRM. Flux rates were primarily driven by gravimetric moisture, but response was also modulated by other factors, like litter type. Finer portions of litter reached higher wetness and exhibited higher flux, compared to coarse tillers under the same conditions, corroborating previous laboratory measurements (Jacobson and others 2015). Differences in moisture absorbance are likely due to differences in surface area to volume ratio or to physical properties; for example, the waxy cuticle on coarse stems resists moisture uptake, while fine litter absorbs it readily. High proportions of fine litter could thus cause NRM to have a greater impact on decomposition. In the Namib, fine litter constituted roughly 50% of *S. sabulicola* standing litter (unpublished data), but this proportion could be higher in systems dominated by annual grasses. Substrate has been known to be have a strong influence on dew formation (Beysens 1995), and early studies recognized that litter type influenced the relative humidity at which litter becomes wet (Bartholomew and Norman 1947), but physical properties are an under-recognized driver of decomposition (compared to chemical properties, e.g. C:N), and may be especially important for decay of standing litter under NRM. In general, linking standard meteorological descriptors of NRM to litter moisture content deserves further attention; variation in flux rates under NRM (e.g. **Fig 4A**), due to variation in moisture content, was the main contributor to uncertainty in our mass loss estimates (**Fig. 7**, **Fig. S4**), not estimates of NRM frequency.

Going forward, NRM event duration (e.g. number of wet hours) will also be an essential variable for estimating the contribution of NRM to decomposition at any site. Unlike rainfall-induced activity, NRM-induced wetness is not easily captured by water amount. Dawson and Goldsmith (2018) recently estimated the contributions of rain to leaf wetness, and Beysens (2016) highlights opportunities to estimate dew from meteorological data, but quantifications of wet periods stimulated by all forms of NRM (fog, dew, and high relative humidity) are lacking. We found that leaf wetness sensors recorded most NRM events, but may underestimate NRM decomposition because litter can stay wet (and respiring) for longer than sensors remain wet. So although we were able to estimate leaf wetness relatively accurately using a likelihood function, refinements are possible. We suggest any effort to quantify decomposition-relevant NRM at a site would start with simultaneous measurements of hourly RH, leaf wetness (each at the height of the litter of interest (Sentelhas and others 2008)), and litter gravimetric moisture (potentially taking advantage of novel methods (Wang and others 2015)). These data could serve to calibrate estimates of NRM to include those likely to induce decomposition, and also to estimate wet hours from leaf wetness or RH in past (or to-be-collected) standard meteorological data. With no previous knowledge of these relationships at a site, our data suggest that assuming wet hours occur above a threshold of 85% RH (Dix and Webster 1995) can be a good starting point for estimating NRM.

We found that NRM events also correspond to particular meteorological conditions that may need to be accounted for as we determine the cumulative contribution of these periods to annual mass loss. For instance, NRM occurs at lower temperatures than rain events in dry sites (**Table 2**), in line with the relatively lower water holding capacity of cooler air. Previous investigations of microbes in drylands focus on traits allowing survival at extremely high temperatures (Sterflinger and others 2012), but many of these organisms have broad thermal optima (e.g. (Sterflinger and others 2012; Jacobson and others 2015)), and may actually be more active during cool moist NRM periods (Jacobson and others 2015). From a modeling perspective, even though NRM decomposition might respond to temperature and moisture in a similar way to rainfall-mediated decomposition, because NRM consistently occurs at cooler temperatures, it might induce lower hourly C flux. Future studies of microbial traits that influence rain- and NRM-decomposition should examine activity at temperatures relevant to these events, rather than the thermal extremes during which microbes are desiccated and inactive.

### Broader role of NRM in ecosystems

The ecological effects of NRM decomposition could extend far beyond decomposition of surface litter during NRM periods, as we documented here. In drylands, nighttime NRM may be a key component that alternates with daytime photodegradation to induce greater decomposition than these processes contribute individually (Almagro and others 2015; Gliksman and others 2017; Lin and others 2018). NRM and UV-PAR can also contribute to surface priming in standing litter (Wang and others 2017a), and the resulting leaching of DOC can contribute to soil carbon dynamics (Campbell and others 2016). Finally, we previously showed that NRM decomposition increased surface nitrogen content in grass litter 2-fold, and that termites preferentially consumed this litter (Jacobson and others 2015). Termites and other detritivores are essential prey for higher trophic levels in most arid ecosystems (Crawford and Seely 1995). The importance of NRM-mediated decomposition may cascade through trophic levels independent of the effects of rainfall on subsurface decomposition.

Even more broadly, additional studies are needed to understand the differential effect of NRM on carbon sources and sinks, particularly in grasslands, where surface litter may comprise more than two-thirds of annual net primary production (Polis 1991). In addition to litter decomposer communities, NRM can also stimulate surface soil crusts, lichen fields, and hypoliths (Wang and others 2017b), plant growth (Dawson and Goldsmith 2018), and soil microbial activity (Carbone and others 2011). In the Namib, NRM stimulates the growth of perennial bunch grasses as it drips from aboveground structures to shallow roots (Ebner and others 2011), and nutrients leached via these moisture droplets could be recycled to growing plant material and contribute to nutrient islands (e.g. (Abrams and others 1997)). NRM may also influence these processes as it alters the timing of moisture availability, an important regulator of biogeochemical dynamics in grasslands (Jacobson and Jacobson 1998; Austin and others 2004; Borken and Matzner 2009), but one in which NRM is not currently considered.

Accurately predicting carbon dynamics worldwide relies on an improved understanding of the drivers of decomposition processes. We demonstrated that NRM is an important component of decomposition of surface litter in hyper-arid and mesic grasslands, and our first effort to model NRM highlights the complexities involved in using this component to improve mass loss predictions. In future decades, the frequency and duration of fog, dew, and RH are predicted to shift (Haensler and others 2011; Engelbrecht and others 2015; Tomaszkiewicz and others 2016), and may already be changing. Takle (2011) reports that Iowa has experienced an increase in summer dew-point temperature over the last several decades, yielding an increase in atmospheric water vapor over the period. Additional monitoring is needed to assess shifts in NRM. Notably, changes in fog and dew patterns may be distinct from one another (e.g. in the Namib (Kaseke and others 2017; Wang and others 2017b)), and from shifts in rainfall. Our study shows – with empirical evidence and extrapolation – that shifts in both rain and NRM will need to be accounted for to accurately predict future decomposition rates.

## Acknowledgements

We thank Robert Logan for insightful discussions and field assistance, and Robert and Esbeiry Cordova-Ortiz for helpful comments on the manuscript. We also thank Gobabeb Research and Training Centre and Roland Vogt for meteorological insight and data acquisition. We thank two anonymous reviewers in a previous submission of this manuscript to Ecosystems. Samples were collected under Namibia Ministry of Environment and Tourism research/collecting permit number 2228/2016. PJ and KJ received funding from Grinnell College, and SE from Michigan State University, and National Geographic Society (WW440-16 to SE). KTB is grateful for the support of the Linus Pauling Distinguished Postdoctoral Fellowship program, part of the Laboratory Directed Research and Development Program at Pacific Northwest National Laboratory, a multiprogram national laboratory operated by Battelle for the U.S. Department of Energy. This is Kellogg Biological Station contribution number 2128.

