## Supplemental Material for "Non-rainfall moisture: a key driver of carbon flux from standing litter in arid, semiarid, and mesic grasslands"

**Supplemental methods**

*Justification for litter rack method*

Although litter bags are the standard method for assessing decomposition dynamics, they are known to alter litter microclimate (Bokhorst and Wardle 2013). We suspected litter bags would alter NRM, so we compared moisture content of litter in standard 1-mm nylon mesh litter bags to litter unenclosed (resting on a 10-cm square piece of 1-mm nylon screen on the ground surface) under two dew events in Iowa, and found that litter in bags had significantly lower moisture content (22-34% less) than litter unenclosed, depending on the duration of the dew event (Table S1). In addition, we suspected that litter bags could not easily mimic the vertical position or height of natural standing litter, where relative humidity and temperature regimes differ from the surface (Wang and others 2017; Gliksman and others 2018). We found that in both Namib sites, tillers in traditional bags had lower annual mass loss than thoser in open racks (Table S1). Thus, we used litter racks to measure mass loss and respiration of standing litter under NRM. Racks were placed in a vertical position, at the height of the litter of interest, and had no barriers (e.g. screens) to alter NRM exposure.

**Supplementary Tables**

**Table S1**. Tests of the effect of containment method on gravimetric moisture under NRM in Iowa

| Response  Variable | Site | Method | Mean  (%) | SD | SE | N | t-test*  p-value |
| --- | --- | --- | --- | --- | --- | --- | --- |
| Gravimetric moisture (%) | Iowa | Bag | 61.33 | 6.43 | 3.7 | 3 | 0.05 |
|  |  | Screen | 95.7 | 13.4 | 7.8 | 3 |  |
| Gravimetric moisture (%) | Iowa | Bag | 25.33 | 2.08 | 1.2 | 3 | 0.014 |
|  |  | Screen | 47.67 | 4.04 | 2.3 | 3 |  |
| Mass loss (%) | Namib E | Litterbag | 1.820 | 0.628 | 0.28 | 5 | <0.001 |
|  |  | Rack | 4.58 | 1.40 | 0.44 | 10 |  |
| Mass loss (%) | Namib W | Litterbag | 1.24 | 2.27 | 1.0 | 5 | 0.002 |
|  |  | Rack | 4.85 | 2.22 | 0.70 | 10 |  |

*testing null hypothesis of no difference between containment method

**Table S2.** Statistical tests of the difference between temperature and relative humidity of NRM-hours and rain-hours, corresponding to Table 2.

| Variable | Site | Variances | Test | Mean NRM | N | Mean Rain | N | p-value |
| --- | --- | --- | --- | --- | --- | --- | --- | --- |
| Temperature | Sevilleta | equal | t-test | 13.8±11.2 | 40986 | 12.6±8.6 | 882 | 2.14 x 10^-5^ |
|  | Namib E | unequal | Wilcox | 12.5 ±6.6 | 414 | 21.5 ±5.3 | 49 | 8.68 x 10^-14^ |
|  | Namib W | unequal | Wilcox | 12.9 4.0 | 1508 | 18.2 ±2.5 | 14 | 3.44 x 10^-6^ |
|  | Iowa | equal | t-test | 12.8 ±10.1 | 4918 | 13.0 ±7.6 | 347 | 7.00 x 10^-3^ |
| Relative humidity | Sevilleta | unequal | Wilcox | 39.9±23.8 | 40986 | 76.9±17.4 | 882 | 2.20 x 10^-16^ |
|  | Namib E | unequal | Wilcox | 81.1 ±11.6 | 414 | 68.0 ±19.9 | 49 | 6.12 x 10^-7^ |
|  | Namib W | equal | t-test | 87.7 ±9.0 | 1508 | 72.8 ±8.8 | 14 | 7.28 x 10^-4^ |
|  | Iowa | unequal | Wilcox | 93.5 ±7.0 | 4918 | 97.9 ±5.9 | 347 | 2.20 x 10^-16^ |

**Table S3**. Summary of all NRM events under which respiration flux measurements were taken.

| **Event**  **Type** | **Site** | **Time of day** | **Rel Hum (%)** | **Temp**  (°**C)** | **Wetness (%)** | **Flux^+^** | **Flux SD^+^** | **Grav moisture (%)** | **Grav moisture SD** |  | **In reg?^++^** |
| --- | --- | --- | --- | --- | --- | --- | --- | --- | --- | --- | --- |
|  |  |  |  |  |  |  |  |  |  | **N** |  |
| **Coarse litter (tillers)** | |  |  |  |  |  |  |  |  |  |  |
| Dew**^§^** | Namib West | 22:08 | 94.8 | 11.1 | 22.9 | 0.24 | 0.19 | 16.92 | 5.20 | 10 | Y |
|  |  | 0:45 | 96.6 | 9.4 | 58.8 | 0.25 | 0.09 | 19.02 | 3.30 | 10 | Y |
|  |  | 5:40 | 90.2 | 8.0 | 57.7 | 0.089 | 0.058 | 17.53 | 2.30 | 9 | Y |
|  |  | 9:15 | 37.7 | 19.4 | 1.8 | 0.081 | 0.051 | 15.29 | 2.97 | 10 | N |
|  |  | 15:00 | 20.2 | 24.9 | 0.0 | 0.088 | 0.055 | 5.01 | 1.06 | 10 | N |
| Dew | Namib West | 6:15 | 70.6 | 17.8 | 0.0 | 0.065 | 0.049 | 4.77 | 4.65 | 10 | N |
| Fog | Namib West | 5:30 | 100.0 | 6.6 | 68.8 | 0.66 | 0.34 | 35.14 | 11.71 | 5 | Y |
|  |  | 11:10 | 44.0 | 15.9 | 0.0 | 0.021 | 0.024 | 10.05 | 1.57 | 5 | N |
| Dew | Namib East | 7:00 | 89.9 | 4.7 | 37.1 | 0.064 | 0.045 | 6.38 | 4.48 | 10 | Y |
| Rain | Namib West | 0:21 | 75.8 | 18.5 | 1.2 | - | - | 9.73 | 5.77 | 10 | N |
|  |  | 7:53 | 70.8 | 19.0 | 11.8 | - | - | 32.80 | 16.34 | 10 | N |
| Fog | Namib West | 6:45 | 89.9 | 16.3 | 27.5 | 0.72 | 0.20 | 18.06 | 2.84 | 10 | Y |
|  |  | 9:22 | 55.5 | 22.0 | 1.8 | 0.02 | 0.02 | 9.73 | 1.49 | 10 | N |
| Fog | Namib West | 6:00 | 98.7 | 16.7 | 53.1 | 0.91 | 0.28 | 30.91 | 4.10 | 10 | Y |
|  |  | 8:13 | 82.6 | 18.3 | 1.8 | 0.80 | 0.37 | 23.11 | 3.40 | 10 | Y |
| Fog | Namib West | 10:15 | 53.2 | 23.1 | 1.8 | 0.19 | 0.11 | 14.10 | 4.59 | 10 | N |
| Dew | Iowa | 21:07 | - | - | - | 0.004 | 0.01 | 1.80 | 2.49 | 5 | Y |
|  |  | 5:45 | - | 7.1 | - | 0.38 | 0.12 | 20.70 | 5.93 | 5 | Y |
| Rain | Iowa | 5:15 | 100.0 | 16.4 | - | 0.68 | 0.23 | 21.06 | 0.82 | 5 | Y |
| Dew | Iowa | 23:35 | 89.0 | 8.9 | Wet | 0.89 | 0.43 | 15.56 | 4.96 | 11 | Y |
|  |  | 6:52 | 92.0 | 9.4 | Wet | 0.51 | 0.27 | 12.53 | 3.56 | 9 | Y |
| **Fine litter** | |  |  |  |  |  |  |  |  |  |  |
| Dew | Namib West | 0:45 | 96.6 | 9.4 | 58.8 | 1.81 | 0.48 | 59.95 | 27.20 | 6 | Y |
|  |  | 6:30 | 96.6 | 8.3 | 58.8 | 1.81 | 0.86 | 67.98 | 27.61 | 6 | Y |
| Fog | Namib West | 7:40 | 100.0 | 7.0 | 73.0 | 2.53 | 0.67 | 145.20 | 41.43 | 5 | Y |
| Dew | Namib East | 7:00 | 90.0 | 6.1 | 35.0 | 1.24 | 0.49 | 33.30 | 21.32 | 5 | Y |
| Rain | Namib West | 0:45 | 65.4 | 19.6 | 17.1 | - | - | 9.96 | 1.61 | 10 | N |
|  |  | 8:15 | 62.5 | 21.0 | 1.2 | - | - | 20.50 | 4.06 | 10 | N |
| Fog | Namib West | 6:53 | 89.9 | 16.3 | 27.5 | 2.41 | 0.63 | 37.79 | 10.19 | 10 | Y |
|  |  | 9:29 | 62.2 | 20.0 | 1.8 | 0.059 | 0.085 | 8.34 | 6.72 | 10 | N |
| Fog | Namib West | 6:05 | 98.3 | 16.7 | 48.0 | 2.63 | 0.41 | 103.09 | 14.75 | 10 | Y |
|  |  | 8:16 | 80.8 | 18.5 | 1.8 | 2.33 | 0.31 | 43.32 | 7.88 | 10 | Y |
|  |  | 9:35 | 58.8 | 21.8 | 1.8 | 0.52 | 0.24 | 12.10 | 5.45 | 10 | N |
| Dew | Iowa | 5:45 | - | 7.1 | - | 1.02 | 0.79 | 113.80 | 26.11 | 5 | Y |
| Dew | Iowa | 22:30 | 89.0 | 8.9 | - | 0.95 | 0.19 | 28.73 | 11.19 | 10 | Y |
|  |  | 6:52 | 92.0 | 9.4 | - | 0.97 | 0.39 | 25.22 | 7.20 | 10 | Y |

**^§^**This event (all time points) shown graphically in Fig. 2

^+^units: mg CO_2_-C/g litter/day; SD=standard deviation

^++^included in regression; Y = Yes, included in regression, N= No, not included in regression because measured during drying phase (see Fig. 2, S3A).

**Table S4.** Statistical tests examining difference in litter flux in nonsterile and sterile tillers to test for biological origin of C flux.

|  |  |  | Gravimetric moisture (%) | | | Flux (mg CO_2_-C/g litter/day) | | |
| --- | --- | --- | --- | --- | --- | --- | --- | --- |
| Event type | Time of day | Litter type | Mean (SE) | T-value | p-value | Mean (SE) | T-value | p-value |
| Dew | 5:45 | Nonsterile | 20.70 (2.7) | 1.37 | 0.213 | 0.85 (0.19) | 3.55 | 0.024 |
|  |  | Sterile | 14.38 (3.8) |  |  | 0.136 (0.054) |  |  |
| Rain | 15:15 | Nonsterile | 21.06 (1.8) | 3.57 | 0.016 | 1.418 (0.14) | 6.55 | <0.01 |
|  |  | Sterile | 35.62 (3.6) |  |  | 0.326 (0.088) |  |  |

**Supplementary Figures**

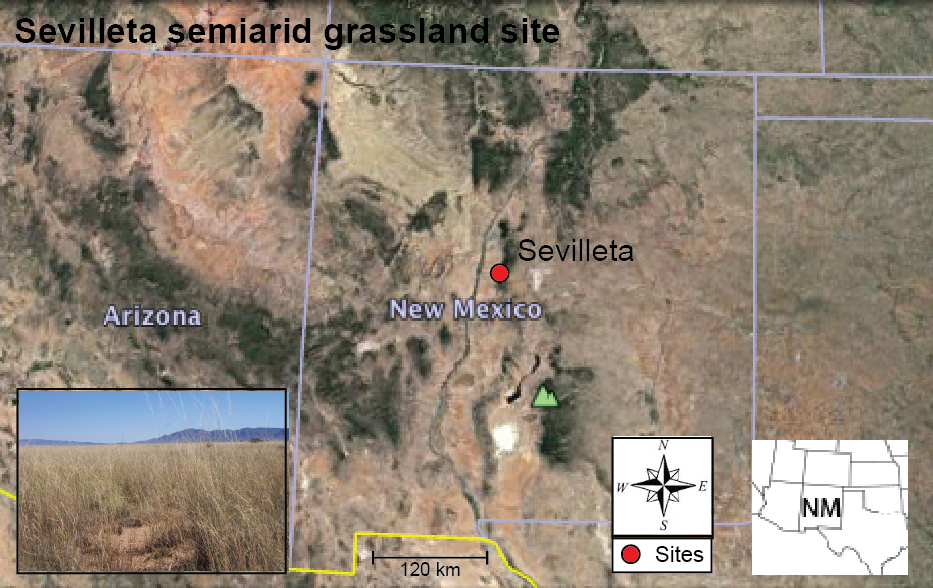

**Figure S1**. Sevilleta, New Mexico site. We used the Sevilleta site to test the robustness of our estimate of NRM decomposition at a semi-arid grassland. . We did not collect litter decomposition, carbon flux or leaf wetness sensor data at this site, but rather estimated NRM using relative humidity from long-term meteorological data, extrapolated potential C loss due to rain and NRM, and compared results with published mass loss data.Map indicates Deep well site at Sevilleta National Wildlife Refuge where RH and mass loss data were collected. Inset shows semi-arid grassland dominated by *Bouteloua eriopoda* and *B. gracilis*

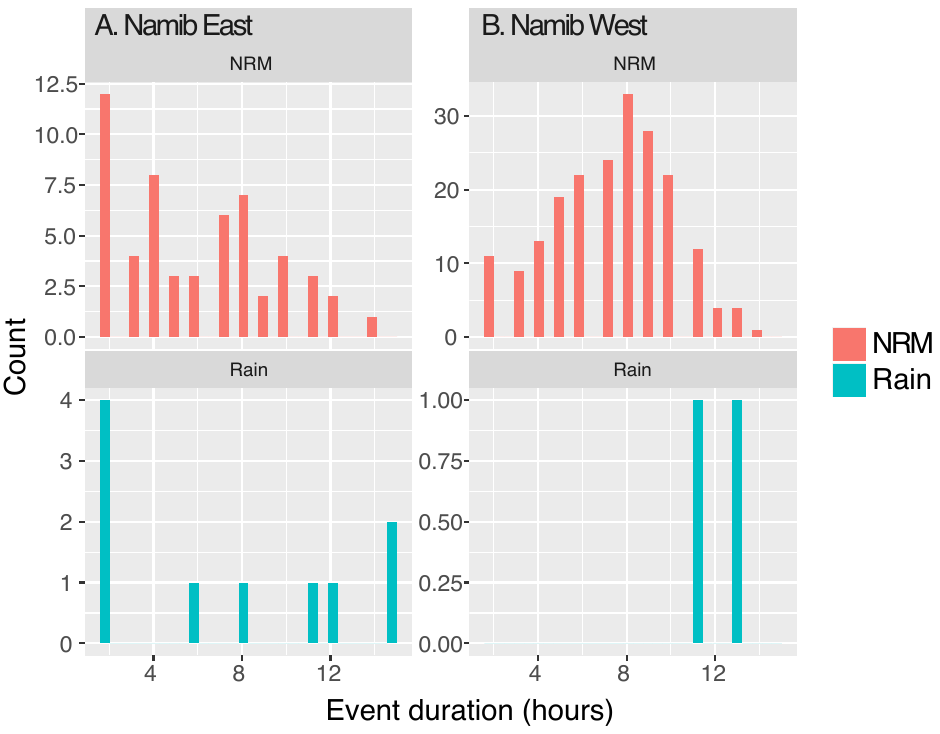

**Figure S2**. Histograms showing the duration (in hours) of a single continuous rain or NRM event in the Namib. Mean NRM duration was significantly higher in Namib West (mean 7.3 h, N=202 NRM events) than Namib East (mean 6.0 h, N=55 NRM events) (t-test, p=0.007).

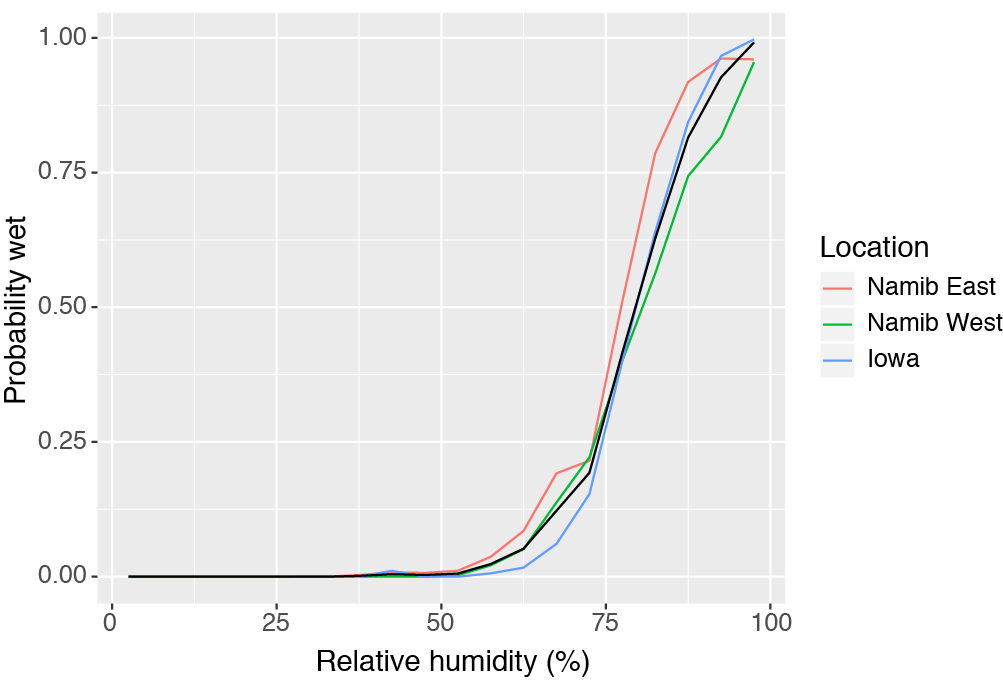

**Figure S3.** Likelihood of “wet” leaf wetness sensor for a given RH, excluding rain events. Based on the global likelihood function (mean across sites, black line), we created a weighted sum for number of hours in each site that the sensor was wet due to NRM.

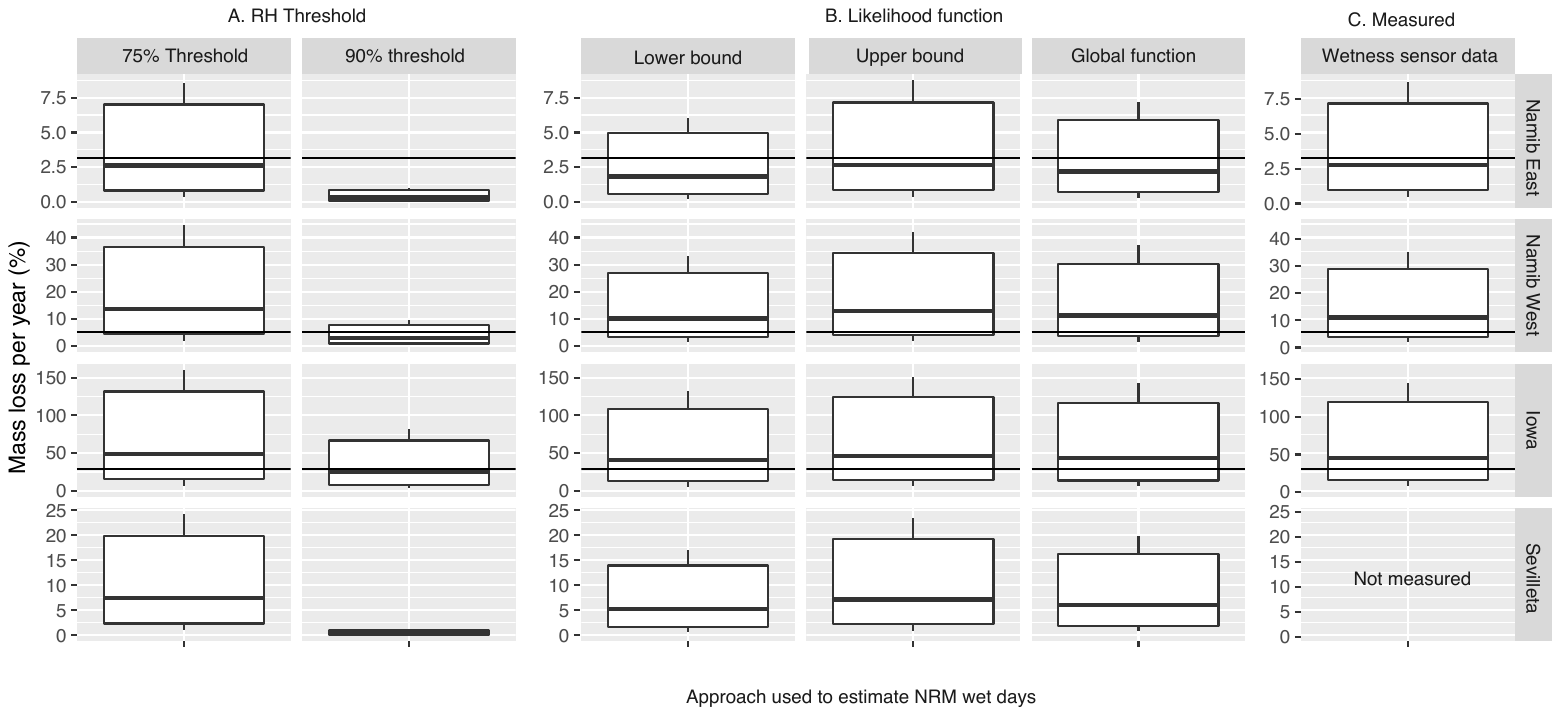
**Figure S4**. Estimates of annual mass loss per year, showing sources of uncertainty generated from respiration extrapolation (variation within plot), RH threshold used (left two columns), and likelihood function used. Global likelihood is shown in Fig. 6 in the main text. The majority of the uncertainty in our estimates in Fig. 6 resulted from the wide range of flux measurements that occur under NRM, but use of RH threshold to estimate NRM frequency, without site-specific calibration using leaf wetness sensors, could also be a source of uncertainty. Black line in Namib and Iowa sites indicates empirically-measured mass loss (mean, standardized to annual scale).
